# Higher stress and immunity responses are associated with higher mortality in reef-building coral exposed to a bacterial challenge

**DOI:** 10.1101/059444

**Authors:** Rachel M. Wright, Carly D. Kenkel, Carly E. Dunn, Erin N. Shilling, Line K. Bay, Mikhail V. Matz

## Abstract

Understanding the drivers of intraspecific variation in susceptibility is essential to manage increasingly frequent coral disease outbreaks. We challenged replicate fragments of eight *Acropora millepora* genotypes with *Vibrio* spp. to quantify variation in lesion development and to identify host and coral-associated microbial community properties associated with resistance. While *Vibrio* spp. remained relatively rare in the microbiome of challenged corals, other stress-associated microbial taxa significantly increased in abundance. Contrary to expectations, higher constitutive immunity and more active immune responses did not confer higher resistance to bacterial challenge. Furthermore, more pronounced gene expression responses to bacterial challenge were associated with higher rather than lower mortality. A newly developed gene expression assay based on two genes related to inflammation and immune responses, *deleted in malignant brain tumors 1* and a matrix metalloproteinase, predicted mortality under *Vibrio* treatment both in the initial experiment and in a validation experiment involving another 20 *A. millepora* genotypes. Instead of mounting more robust responses, resistant corals were largely unaffected by the bacterial challenge and maintained gene expression signatures of healthier condition, including elevated fluorescent proteins and ribosomal biosynthesis along with diminished ubiquitination. Overall, our results support the view that coral disease and mortality is commonly due to opportunistic pathogens exploiting physiologically compromised hosts rather than specific infections, and show, contrary to the prevailing wisdom, that greater immune responses do not necessarily translate into greater disease resistance.

## Significance Statement

Despite its disastrous impact on reefs globally, many aspects of coral disease remain unclear. We find that susceptibility to bacterial challenge in an Indo-Pacific coral, *Acropora millepora,* is associated with increased constitutive stress levels and, surprisingly, with more pronounced immune and gene expression responses to the bacteria. These results and additional microbiome characterization support the view that coral disease is caused by compromised physiological status leading to infection by omnipresent opportunists, explaining why few etiological agents of coral disease have been confidently identified and highlighting the role of abiotic stressors in promoting coral disease outbreaks.

## Introduction

Global declines in coral cover are compounded by a variety of diseases (1, 2), many of which are ambiguously defined by macroscopic characterizations of lesions (1–3). Several bacterial species from the genus *Vibrio* have been implicated as etiological agents of some coral diseases (4–6), but these bacteria may act merely as opportunistic pathogens exploiting compromised hosts (7). Indeed, host immune health is considered to be a major determinant of disease transmission dynamics (8, 9). Corals, like all invertebrates, rely entirely on innate immunity for protection from invading pathogens. Features of innate immunity in corals include molecular pattern recognition (10), secreted antimicrobial macromolecules (11), cellular responses *(e.g.,* phagocytosis) (12, 13), and physical barriers *(e.g.,* mucus) (14, 15). Melanin deposits serve as another physical barrier against invading pathogens (16). The melanin synthesis cascade is activated when pathogen recognition triggers cleavage of prophenoloxidase (PPO) to phenoloxidase (PO). Reactive oxygen species (ROS) produced during melanin synthesis contribute to its cytotoxic effects on pathogens, but also cause self-harm (17) that must be countered by antioxidant enzymes such as catalase (CAT) and peroxidase (POX).

Field surveys of naturally occurring coral disease events show variability in disease outcomes among conspecifics, despite the fact that the colonies are exposed to the same environmental stressors and, presumably, the same potential pathogens (8, 18). One possible explanation is that some corals resist disease by making greater contributions to constitutive immunity and coral taxa investing more in innate immunity (*e.g.*, production of cytotoxic defenses) are less likely to suffer infectious disease outbreaks (19, 20). However, no laboratory experiments have yet confirmed these differences in susceptibility at the intraspecific level or investigated the molecular basis of differential disease outcomes among individuals.

In this study, we comprehensively examine coral host immune activity, genome-wide gene expression, *Symbiodinium* profiles, and coral-associated microbial communities to understand the physiological features underpinning disease resistance in a reef-building coral. Three fragments of eight colonies of *Acropora millepora* from two populations on the Great Barrier Reef (GBR) were individually challenged in a full-factorial design with putative bacterial pathogens *(Vibrio owensii* and *V. diazotrophicus)* and mechanical abrasion and monitored over a time-course of lesion development and progression. Unchallenged fragments were used for baseline measures to correlate constitutive health parameters with survivorship, while postchallenge samples were analyzed for colony-specific responses to the bacterial challenges.

## Results

***A. millepora* genotypes show significant differences in mortality.** Survival analysis revealed significant differences in mortality among individuals (Fig. 1A, p < 0.001). One genotype, W30, experienced no mortality throughout the experiment. Overall, corals from Lizard Island experienced significantly higher mortality than corals from Wilkie Island (p < 0.001). Bacterial treatment increased mortality (control vs. treated p < 0.001), but there was no significant difference in mortality between corals challenged with *V. owensii* or *V. diazotrophicus* (Fig. S1A, p = 0.108). Similarly, abrasion did not significantly affect survival (Fig. S1B, p = 0.16). No significant differences in mortality were detectable between corals treated with either bacteria or between abraded and non-abraded corals, therefore for subsequent analyses all the samples were regarded as just two experimental groups: bacteria-treated (N = 12 per genotype) or control (N = 6 per genotype).

**Higher constitutive immune activities and responses do not correlate with higher survival.** PO and PPO activities were measured as a proxy for cytotoxic defenses via the melanin-synthesis pathway and antioxidant capabilities were assessed by measuring CAT and POX activities in replicate fragments of all genotypes from samples taken at the conclusion of the challenge experiment. The active form of the melanin-forming enzyme, PO, was significantly elevated in bacteria-treated corals compared to controls (Fig. 1B; p = 0.04). No differences with respect to treatment were observed for the inactive form of phenoloxidase (PPO, p = 0.24). Antioxidant activities were significantly increased (CAT: p = 0.008 and POX: p = 0.03) in treated corals relative to control fragments (Fig. 1C-D). While the bacteria-treated corals were clearly launching an immune response, neither the magnitude of these immune responses nor higher constitutive levels were associated with higher survival rates. On the contrary, CAT (p = 0. 006), POX (p < 0.001), and PO (p = 0.04) activities were higher in corals that experienced higher mortality. However, these relationships were largely driven by a single genotype, W30, that did not develop lesions throughout the experiment. No significant association between enzymatic activities and survival remained when this individual was removed from the analysis.

**Fig. 1.**
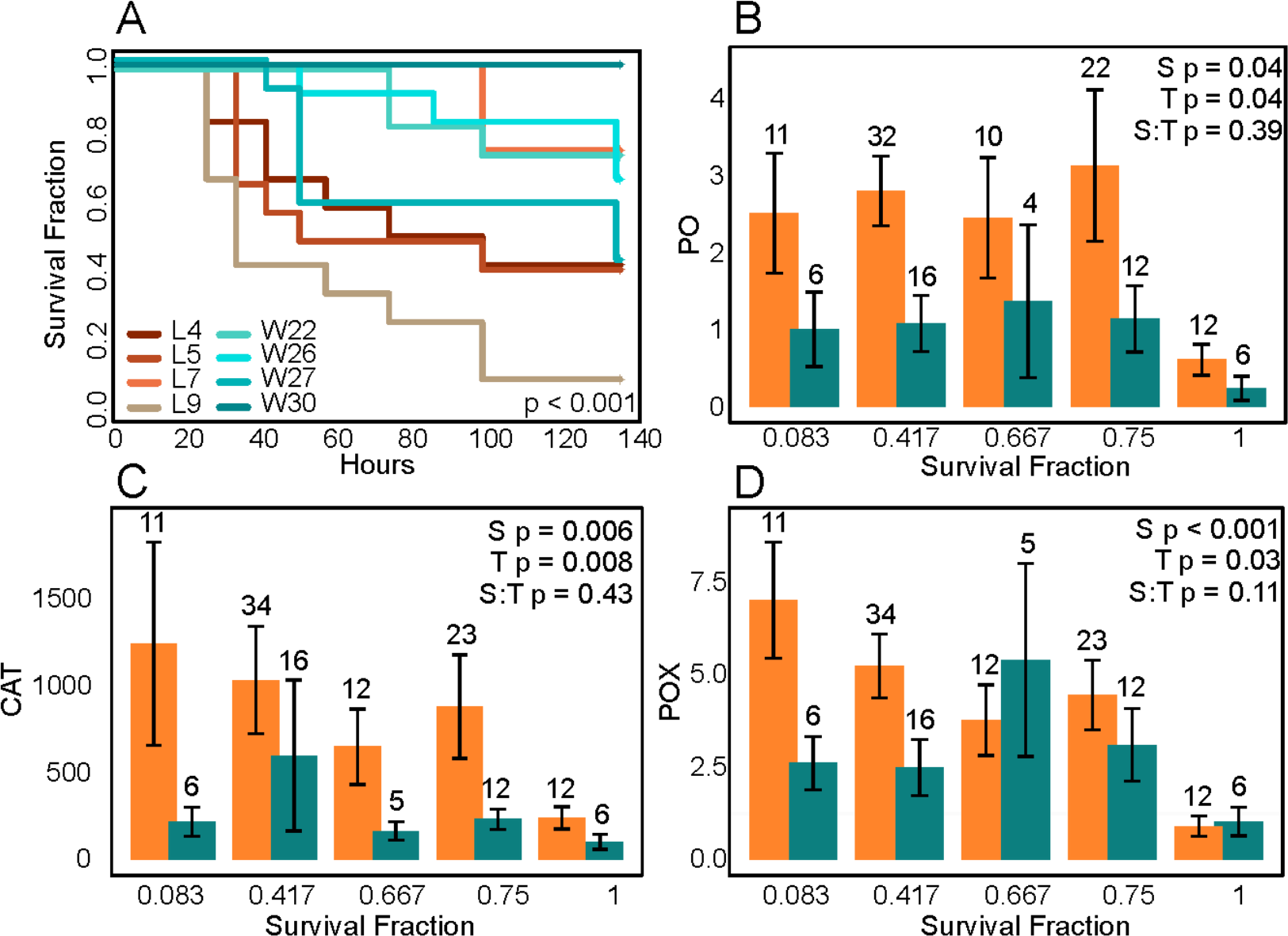
*A. millepora* survival and immune activities after bacterial challenge. (A) Survival by genotype. P-values were generated by Cox proportional hazards models testing the effect of genotype. L4-L9 = corals from Lizard island. W22-W30 = corals from Wilkie Island. (B-D) Mean (±SEM) phenoloxidase (B, PO), catalase (C, CAT), and peroxidase (D, POX) activities are represented as Δ absorbance mg protein^-1^ min^−1^. P-values were generated by generalized linear mixed models testing the effects of survival (S), treatment (T), and their interaction (S:T) with a random effect of genotype. Numbers above bars indicate sample size.

**Baseline gene expression profiles predict survival and lesion development.** Models testing the effect of survival on baseline gene expression yielded 833 differentially expressed genes (DEGs) at FDR = 0.1 Notable among the top DEGs predictive of high survival (FDR = 1e-4) were cyan and green fluorescent proteins and ribosomal proteins (Fig. S2). A rank-based gene ontology analysis revealed significant enrichment of biological processes including elevated oxidative stress defenses and ribosome biogenesis and diminished ubiquitination in highly resistant corals (Fig. 2A).

**Survival-specific gene expression in response to bacterial challenge predicts lesion onset.** A generalized linear model with contrasts between treatments (control vs. bacteria challenge) detected 606 DEGs at FDR = 0.1. Models testing for associations between gene expression and survival yielded 2880 DEGs at FDR = 0.1. Tests of the interaction between survival and treatment yielded three DEGs at FDR = 0.1. Bacterial challenge triggered increased expression of phosphoenolpyruvate carboxykinase, several matrix metalloproteinases (MMPs), and a metalloproteinase inhibitor (Fig. S3A). Genes encoding proteins involved in immunity (interferon gamma) and programmed cell death (apoptosis regulator Bcl-W) were upregulated in bacteria-challenged corals, while deleted in malignant brain tumors protein 1 (*dmbt1*) and a cryptochrome were downregulated. Gene ontology analysis of genes differentially expressed by bacterial treatment found only one significantly enriched term (FDR = 0.1), “small molecule metabolic process”, which was downregulated in treated corals. Notably, the gene expression response to bacterial challenge was predominantly driven by more susceptible genotypes, as expression profiles of the most resistant corals remained similar to the control condition (Fig. 2B). Indeed, our model predicted diminished fold-changes in resistant corals for 97% of all bacteria-related DEGs (Fig. S4). Similarly to the pre-challenge condition, more resistant corals continued to express higher levels of fluorescent proteins (cyan and green) and ribosomal proteins after bacterial challenge, while more susceptible corals exhibited higher expression levels of a stress-activated MAP kinase-interacting protein, ubiquitin ligase, C-type lectin, and complement factor B (Fig. S3B).

**Fig. 2.**
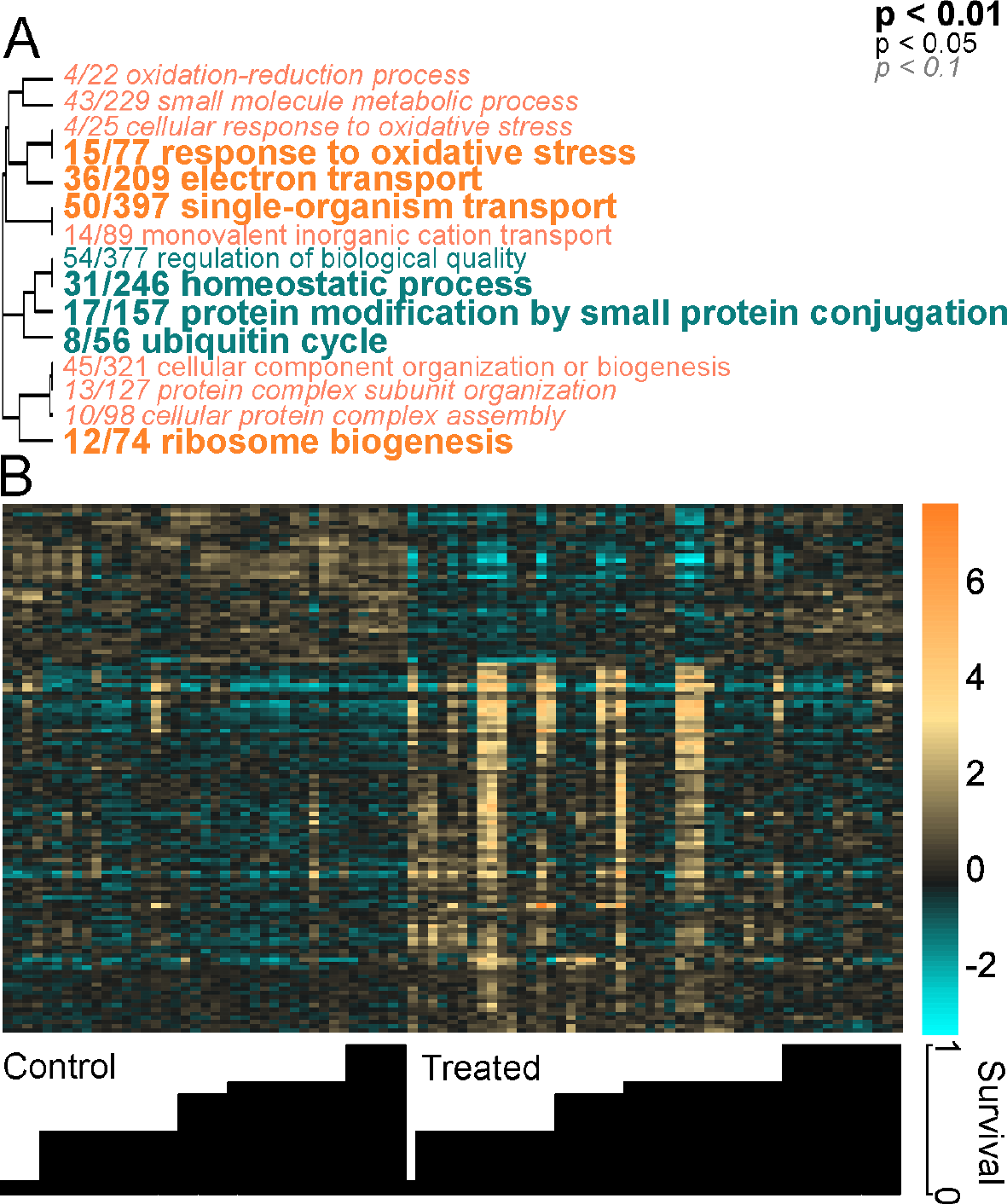
Summary of baseline and response gene expression differences by survival. (A) Biological processes enriched by survival in baseline gene expression profiles. The text color indicates the direction of expression difference between corals with high and low survival (turquoise = elevated in high survival, orange = diminished in high survival). The text size indicates the significance of the term as indicated by the inset key. The fraction preceding the term indicates the number of genes within the term that pass an unadjusted p-value threshold of 0.05. Trees indicate gene sharing among gene ontology categories (categories with no branch length between them are subsets of each other). (B) Heatmap comparing the top DEGs (FDR = 0.1) from response expression profiles by survival fraction. Rows are genes and columns are samples. The color scale is in log2 (fold change relative to the gene’s mean). Genes are hierarchically clustered based on Pearson’s correlations of expression across samples. Samples are clustered first by treatment (control and bacterial treatment) then by increasing survival from left to right.

***Symbiodinium* profiles differ by reef and survival.** *Symbiodinium* communities were profiled using RNA-seq reads mapping uniquely to clade A, B, C, or D *Symbiodinium* transcriptomes. As expected for *A. millepora* in this region of the GBR, clade C dominated most genotypes (6/8, Fig. S5) (21). While corals with the highest survival rate also contained the most D-type symbionts (p < 0.001), corals with the second-lowest survival rate were also significantly enriched with clade D symbionts (p < 0.001).

**Microbial community profiles differ between individuals and in response to bacterial challenge.** Clustering at 97% similarity identified 1238 operational taxonomic units (OTUs). Principal coordinate analysis (PCoA) of weighted UniFrac distances revealed significant differences depending on survival (Fig. 3A). As expected, asymptomatic corals treated with *Vibrio* spp. had higher abundances of *Vibrio* than untreated controls (Table S1; FDR = 0.018). Stress-associated taxa (families) were more abundant in treated corals (p = 0.04, Fig. 3B, Table S2). Notably, some corals with high survival already had the highest proportion of stress-associated taxa in the control treatment (p = 0.014; Fig. 3B). In addition, *A. millepora* with the highest survival rate (genotype W30) had significantly more chloroplast-derived OTUs than other genotypes (Fig. S6B; FDR = 0.001). These taxa likely indicate the presence of endolithic photoautotrophs, corroborated by a visually greener skeleton of W30 than other genotypes (Fig. S6A). Chloroplast-derived OTUs were excluded from subsequent analysis of the bacterial populations.

**Fig. 3.**
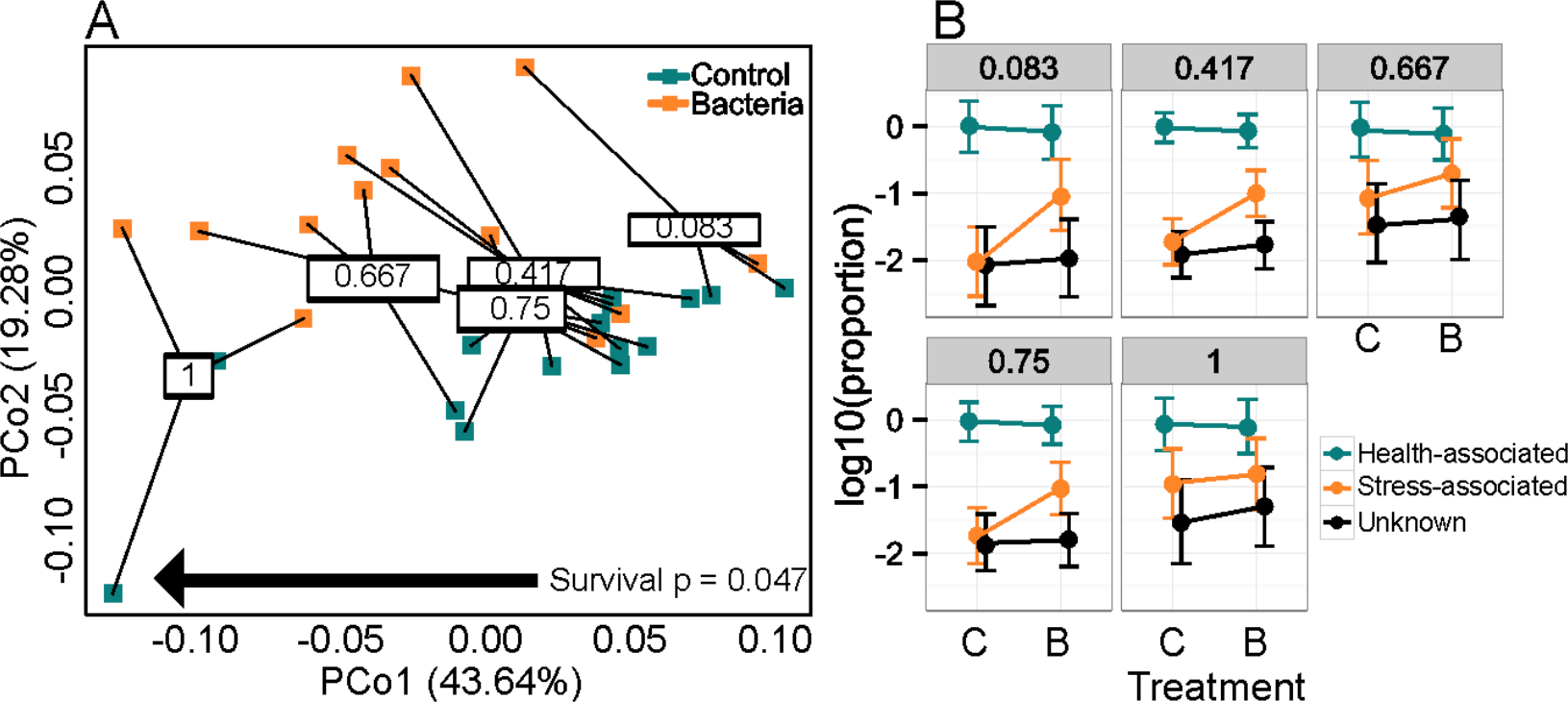
Microbial community composition by treatment and survival. (A) Principal coordinate analysis of microbial community profiles using weighted UniFrac distances. P-values were generated by a PERMANOVA testing the effect of survival fraction. (B) Differences in microbial community composition by treatment (C = control; B = bacteria-challenged) and survival fraction (numbers on top of panels). OTUs were classified as stress-associated or health-associated based on published studies. The plots show posterior means and 95% credible intervals.

**Diagnostic gene expression biomarker identification and validation.** Two candidate survival genes were selected based on their high and dynamic expression: deleted in malignant brain tumors protein 1 (*dmbt1*) and gelatinase and matrix metalloproteinase (*mmp*). *Dmbt1* expression was positively and *mmp* expression was negatively correlated with survival (p < 0.001 for both) (Fig. S7). This gene pair was used in a self-normalizing double-gene qPCR assay, *sensu* (22). The result of the assay was positive for resistant corals and negative for susceptible corals (Fig. 4A). The same relationship held in an independent bacterial challenge experiment performed on an additional 20 *A. millepora* genotypes (Fig. 4B).

**Fig. 4.**
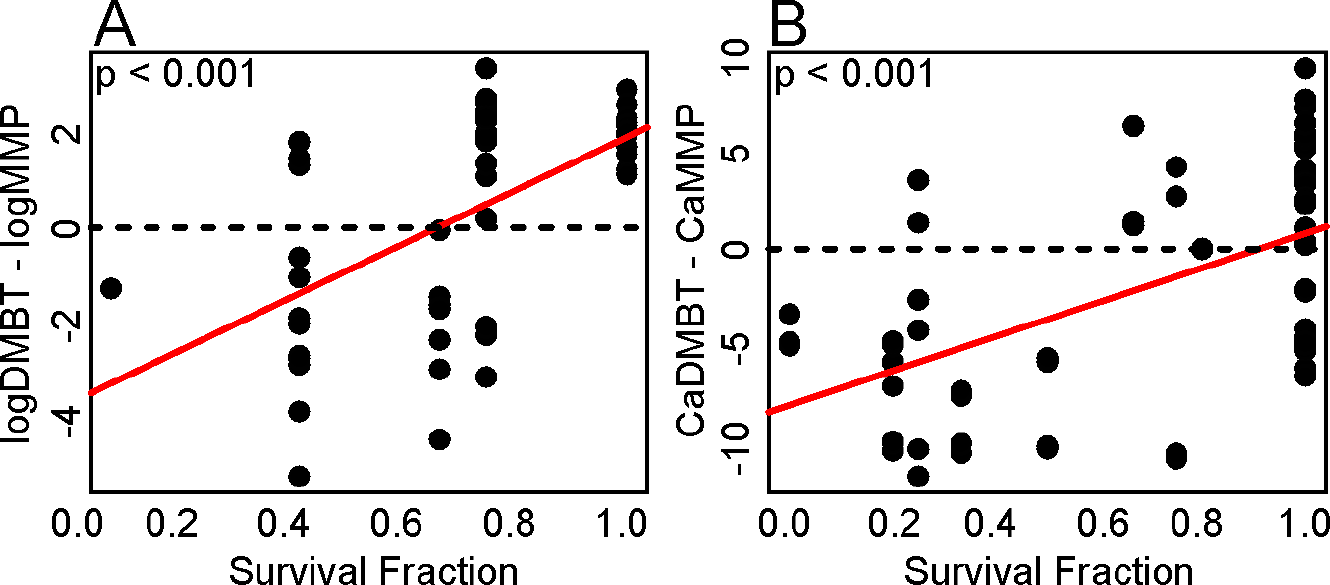
Diagnostic double-gene expression biomarker validation. (A) The difference in RNA-seq derived expression values (log-normalized) of *dmbt* and *mmp* corresponded with survival in the original experiment. (B) The difference in qPCR-derived Ca values for *dmbt* and *mmp* also significantly correlated with survival in the validation experiment (n = 20 genotypes). P-values indicate the results of linear models testing the effect of survival on double-gene expression differences. Dotted lines indicate double-gene values = 0, below which survival outcomes are unfavorable.

## Discussion

**Higher immune response to bacterial challenge does not translate into lower mortality.** The current view of coral immunity is that species with higher constitutive and/or inducible immune activity are more resistant to disease (19, 20). However, we found that this hypothesis does not apply to intraspecific differences in disease resistance, as resistance was not correlated with antioxidant and cytotoxic activities, and higher survival was not associated with more robust immune responses to bacterial challenge (Fig. 1B-D). Rather, resistant corals were largely unaffected by the bacterial challenge according to the selected immune activity assays. This unexpected finding suggests that mechanisms other than host immune activity underlie increased resistance to the bacterial challenge.

**Resistant corals show a “healthier” gene expression profile.** Constitutive (baseline) gene expression analysis provided comprehensive insight into the physiology underlying observed differences in predisposition to lesion formation. While we did not observe increased catalase or peroxidase activities in resistant corals, the constitutive upregulation of oxidative stress response genes (Fig. 2A) suggests that these individuals may “frontload” (*sensu* (23)) antioxidant activity in some manner that was not detected by our enzymatic assays. Constitutive frontloading of thermal tolerance transcripts is a proposed mechanism by which some corals outperform conspecifics under heat stress (23), and thus offers a tempting explanation for observed differences in immune performance as well. However, surprisingly few genes involved in stress responses or immunity were upregulated in resistant corals prior to bacterial challenge. Instead, these individuals exhibited a more “healthy” gene expression profile: they had elevated ribosomal components typically associated with higher growth rate (24) as well as fluorescent proteins whose abundances have been linked to health status in corals (25, 26). Another signature of higher resistance was diminished abundance of ubiquitination-related transcripts, *e.g.,* ubiquitin ligases and ubiquitin carboxy-terminal hydrolases. Ubiquitination labels damaged proteins for removal and is a general hallmark of cellular stress (27). Ubiquitin has been shown to be upregulated in heat-stressed corals with high levels of damaged proteins (28–30). Differential expression of ubiquitination-related transcripts suggests some genotypes from both sampling locations may have been experiencing more cellular stress than others at the beginning of the experiment.

**More pronounced gene expression response to *Vibrio* challenge is associated with higher mortality.** Susceptible corals experienced dynamic changes in gene expression in response to bacterial challenge, including upregulated stress response and immune functions (e.g., stress-activated MAP kinase-interacting protein, C-type lectin, ubiquitin ligases, and complement factor B) (Fig. S3). Notably, expression profiles of the most resistant, treated corals were more similar to the expression profiles of untreated individuals (Fig. 2B and Fig S4). Bacteria-treated resistant corals maintained their initially higher expression levels of fluorescent proteins (green and cyan) and ribosomal proteins along with lower ubiquitination (Figs. S2, S4). Taken together with the lack of elevated immune activity in resistant corals (Fig. 1B-D), our conclusion is that resistant corals survived because they were less sensitive overall to the bacterial challenge rather than because they launched a more robust response.

***Symbiodinium* and bacterial profiles characterized in resistant and susceptible corals.** Corals with clade D *Symbiodinium* communities can be more resistant to disease than those hosting clade A symbionts (31), however, no significant association between algal symbiont composition and survivorship was detected here. Clade D dominated colonies were among the most and least resistant to bacterial challenge, highlighting the need for further investigation of the symbiont’s role in the host’s predisposition to disease. *Vibrio* treatment caused an increase in the abundance of stress-associated taxa including Alteromonadaceae, Pseudoalteromonadaceae, Rhodobacteraceae, and of course, Vibrionaceae. Although the proportion of *Vibrio* was elevated in challenged corals (Table S1; FDR = 0.018) it remained low, suggesting that the introduced *Vibrio* may not have established an infection. Instead, it appears that the introduced bacteria triggered a disturbance in the coral-associated microbiome that facilitated the proliferation of resident opportunists. Microbiome shifts favoring potential pathogens were evident in all genotypes except the most resistant (Fig. 3B). Surprisingly, resistant corals tended to harbor elevated levels of these potentially harmful bacteria even before the challenge (Fig. 3B), which drove the association between bacterial composition and survivorship (Fig. 3A). This finding suggests that resistant corals are not more efficient in excluding harmful bacteria but are generally less sensitive to their presence, which aligns well with their insensitivity to bacterial challenge at the gene expression and physiological levels, as discussed above.

The most striking difference in microbial composition is the most resistant coral (W30) harbored more chloroplast-derived OTUs (Fig. S6B), suggesting the presence of endolithic algae or cyanobacteria. It is tempting to speculate that these microbes facilitated host defense by actively secreting antimicrobial compounds, an ability that has been well characterized in cyanobacteria (32, 33). This putative association between chloroplast-derived OTU abundance and lower mortality under bacterial challenge merits further investigation.

**Genes involved in host-microbe interactions predict the onset of disease.** This study developed and validated a double-gene assay (22) predicting whether an asymptomatic coral is likely to develop disease symptoms in the near future (Fig. 4). This assay can be used to assess risks of mortality during coral disease outbreaks, while the nature of genes constituting it provides key insights into potential mechanisms mediating disease susceptibility. Deleted in malignant brain tumors 1 (*dmbt1*) protein is found in the gut mucosa of humans where it acts as a pattern recognition receptor that maintains mucosal homeostasis by inhibiting bacterial invasion and suppressing inflammation (34, 35). Other transcriptomic studies have found that *dmbt1* was downregulated in oysters upon bacterial challenge (36), upregulated in *Orbicella faveolata* after lipopolysaccharide challenge (37), and upregulated in aposymbiotic sponges compared to sponges infected with clade G *Symbiodinium*, suggesting that *dmbt1* may play a role in mediating symbiosis (38). Elevated *dmbt1* in all control fragments and in resistant corals relative to susceptible, bacterial-challenged corals may signify the role of this protein in maintaining symbiotic associations with commensal microbes. The diagnostic gene that was regulated in the opposite direction, a matrix metalloproteinase (*mmp*), belongs to a family of enzymes with a wide range of functions. The upregulation of MMPs in response to parasitic protists in a gorgonian coral (39) and in *A. hyacinthus* affected with White Syndrome-like symptoms (25) suggests an active role of these proteins in the immune response of cnidarians. This study finds that changes in *dmbt1* and *mmp* may represent some of the earliest coral responses to immune challenge, as they are evident even in asymptomatic corals.

**What causes coral disease?** The bacterial challenge experiment in this study was conceived under the hypothesis that the introduced *Vibrio owensii,* but not *V. diazotrophicus,* would act as an infectious agent. The scenario actually observed was radically different: neither bacterial species proliferated within the host, but both treatments triggered the rise of opportunistic pathogens in the coral microbiome and subsequent development of disease lesions in less healthy corals. We are not the first to argue that a coral disease may be an opportunistic infection exploiting a compromised host (7) and many coral diseases are associated with broad shifts in microbial community composition beyond the rise of a single pathogenic agent (40–42). If, as our results suggest, natural coral disease events are indeed driven predominantly by the host’s health status rather than by activities of true infectious agents, environmental factors influencing coral health (e.g., thermal stress or ocean acidification) might play a much larger role in coral disease than is currently believed.

## Materials & Methods

**Bacterial challenge experiment.** Details concerning coral collection, aquarium conditions, and bacterial culturing are outlined in *SI Materials and Methods.* Briefly, four colonies per reef were collected from reefs near Lizard and Wilkie Islands in the Great Barrier Reef and maintained in common garden conditions for approximately one month. Baseline tissue was preserved for gene expression analysis and colonies were fragmented into 18 fragments per colony. Nine of the 18 fragments received two small abrasions with an airgun blast and all fragments were placed into individual 200 mL containers. Liquid cultures of *Vibrio owensii* strain DY05 and *V. diazotrophicus* were prepared daily from single isolates. Liquid cultures were triple-washed in 0. 4 μM-filtered seawater and diluted to a final concentration of 10^7^ CFU · mL^−1^ in the individual coral containers. Of the nine abraded fragments per genotype, three were challenged with *V. owensii,* three were challenged with *V. diazotrophicus,* and three received daily inoculations of filtered seawater (control). The nine non-abraded fragments received the same treatments. Aquarium water was changed daily preceding each bacterial challenge. Fragments were monitored for tissue loss and photographed twice daily throughout the experiment with a Nikon D300 digital camera (Nikon, Tokyo, Japan). Corals were photographed and preserved in liquid nitrogen when tissue loss was visually estimated at 50% or more of the total surface area of the fragment (*e.g.,* Fig. S6A: L8 and W26). All remaining fragments were frozen in liquid nitrogen and stored at −80°C at the end of the experiment.

**Survival analysis.** The time when fragments suffered ~50% tissue loss was recorded for each fragment. Survivorship analyses were performed for each genotype, reef, abrasion treatment, and bacterial challenge using the Kaplan-Meier estimate of the survival function as implemented by *survfit* in the R package *survival* (43). Cox proportional hazards models fit using the *coxph* command in the *survival* package tested the significance of the effect of reef, genotype, or treatment on survival. The analysis was repeated with one modification: corals that received either *V. owensii* or *V. diazotrophicus* were grouped to compare corals that received a bacterial challenge to the control corals that did not receive a bacterial challenge.

**Enzymatic assays.** Coral tissue was removed and proteins were extracted following established procedures (12, 44, 45). Details are outlined in *SI Materials and Methods.* Differences in normalized CAT, POX, PO, and PPO values were evaluated with respect to treatment and survival fraction, with genotype as a random effect, using *MCMCglmm* (46). All analyses were performed in R version 3.1.3 (47).

**Gene expression.** Genome-wide gene expression was analyzed using tag-based RNA-seq (TagSeq) method (48). Details regarding library preparation and gene expression analysis are outlined in *SI Materials & Methods.* Sample outliers were removed using R package *arrayQualityMetrics* (49) and differential gene expression analysis was performed using *DESeq2* (50). P-values for significance of contrasts between reefs of origin, treatments, survival fractions, and the survival by treatment interaction were generated based on Wald statistics. Empirical false discovery rates were calculated using the package *empiricalFDR.DESeq2* (25). Gene ontology (GO) enrichment analysis was performed using the GO_MWU method that uses adaptive clustering of GO categories and Mann-Whitney U tests (51) based on a ranking of signed log p-values (52). Gene expression heatmaps with hierarchical clustering of expression profiles were created with the*pheatmap* package in R (53).

***Symbiodinium* analysis.** Trimmed and quality filtered RNAseq reads were mapped to *Symbiodinium* clade A, B, C, and D transcriptomes with *bowtie2* (54). A custom perl script generated counts from mapped reads and calculated clade fractions. The R package *MCMC.OTU* (55) was used to implement generalized linear mixed model analysis to test for significant differences in clade abundances.

**Microbial community analysis.** Microbial communities were profiled for eight baseline samples, eight *V. diazotrophicus-treated* samples, eight *V. owensii-treated* samples, and eight control samples. Details regarding sequencing and QIIME (Quantitative Insights Into Microbial Ecology) (56) analysis can be found in *SI Materials & Methods.* Principal coordinate analyses and PERMANOVA (*adonis*) were conducted based on weighted UniFrac distances (57) using the R package *vegan* (58). Significant differences in OTU abundances between reefs of origin, treatments, and survival fractions were assessed using a likelihood ratio test as implemented by the G-test with group_significance.py in QIIME. Individual OTUs were designated as “stress-associated”, “health-associated”, or “unknown” based on previous literature (Table S2). Significant differences in abundances of OTU types between treatment and survival classes were assessed using generalized linear mixed model implemented in the *MCMC.OTU* package in R (55).

**Validation experiment.** The bacterial challenge was repeated in March 2013. *A. millepora* (N = 43 genotypes, 5 fragments per genotype) were challenged daily with 10^6^ CFU · mL^−1^ *V. owensii* DY05 as described above. An equal number of control fragments for each genotype were maintained under ambient conditions (26C). Survival was monitored for seven days of daily bacterial challenges. On the final day, ~1cm^2^ tissue samples were preserved in 100% ethanol and stored at-80°C. These corals were collected under the permit number G12/35236.1 by the Great Barrier Reef Marine Park Authority of Australia. Twenty genotypes spanning a range of survival rates were used in the qPCR validation.

**Quantitative real-time PCR (qPCR) validation of putative biomarkers.** Candidate diagnostic gene expression biomarkers were selected based on differential expression with regard to survival in the response gene expression dataset and had putative functions that could be related to immune defense. Primers (see *SI Materials and Methods*) were designed using Primer3 (59). RNA isolation, cDNA preparation, and qPCR were carried out as previously described (22) with the exception that the RNAqueous Total Isolation Kit (Ambion) was used to isolate total RNA. Linear regression implemented in R was used to test for the relationship between survival fraction and the log-difference in expression between the two candidate genes, as in (22).

## Acknowledgements

We thank David Bourne for providing bacterial cultures and advice. Bioinformatic analyses were completed using Texas Advanced Computing Center resources. This study was supported by grants from NSF (DEB-1054766) to M.V.M. and the Australian Institute of Marine Science to L.K.B. Travel support from the Center for Computational Biology and Bioinformatics at UT Austin was awarded to R.M.W. Two University Co-Operative Undergraduate Research Fellowships were awarded to E.N.S. and C.E.K. Sequencing data have been deposited to the National Center for Biotechnology Information’s Short Reads Archive under accession numbers SRP074065 and SRP073937.

---

**Fig. S1:**
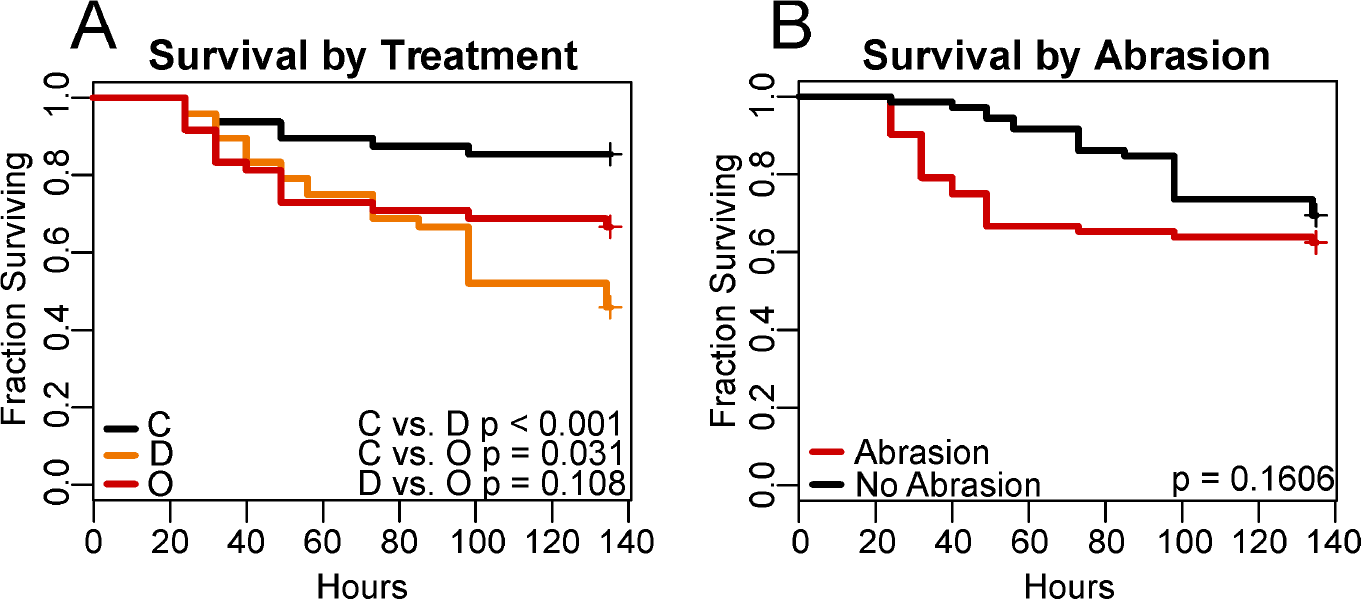
*A. millepora* survival by treatment during bacterial challenge. (**A**) Black, orange, and red lines represent the survival of control, *V. diazotrophicus-challenged,* and *V. owensii-challenged* corals, respectively. (**B**) Black and red lines represent the survival of corals that abraded and non-abraded corals, respectively. P-values were generated by Cox proportional hazards models testing the effect of each treatment.

**Fig. S2:**
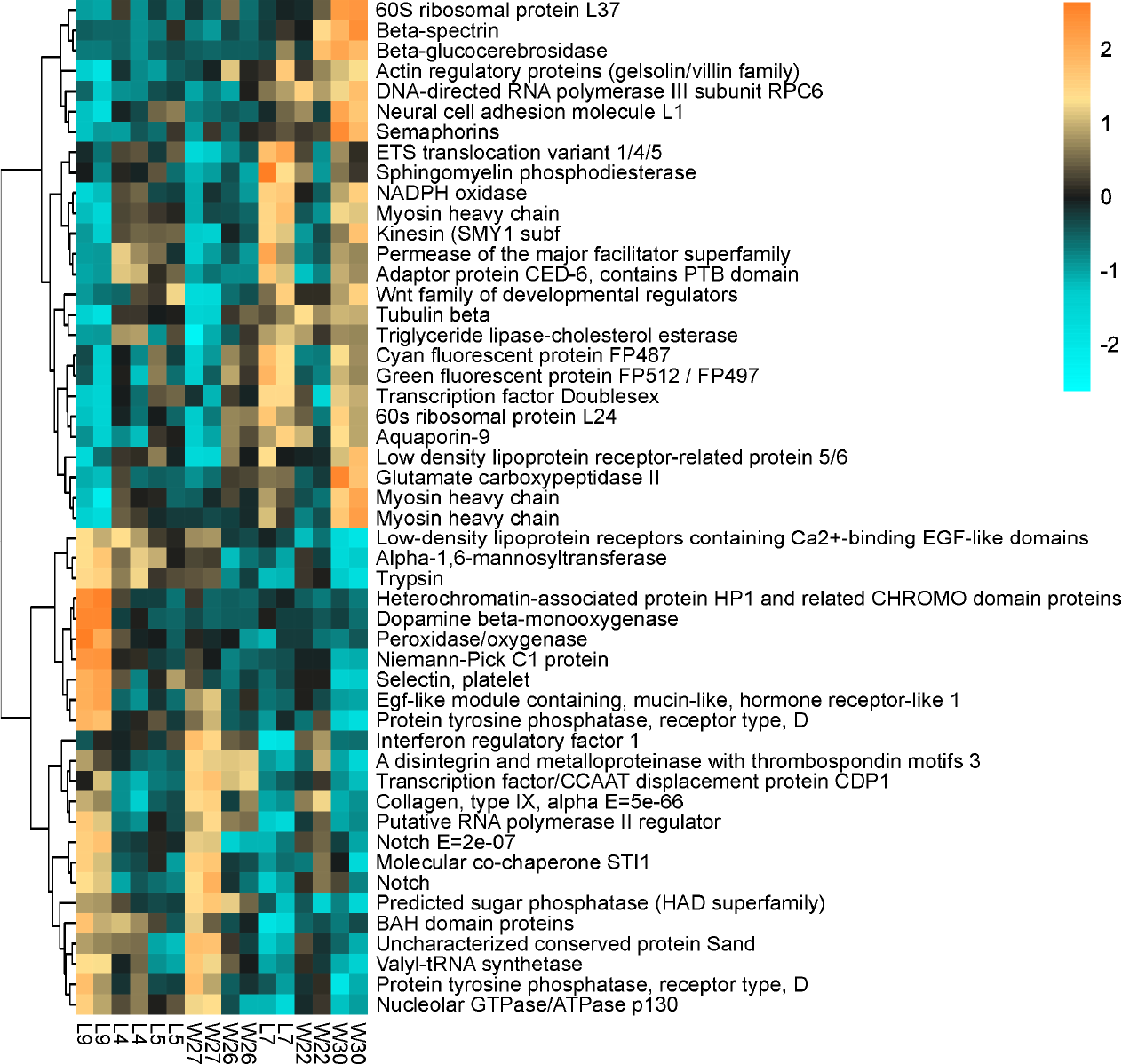
Baseline gene expression heatmap. (Heatmap comparing the top DEGs (FDR = 1e-9) from baseline gene expression profiles by survival fraction. Rows are genes and columns are samples. The color scale is in log2 (fold change relative to the gene’s mean). The tree is a hierarchical clustering of genes and samples based on Pearson’s correlation of expression across samples.

**Fig. S3:**
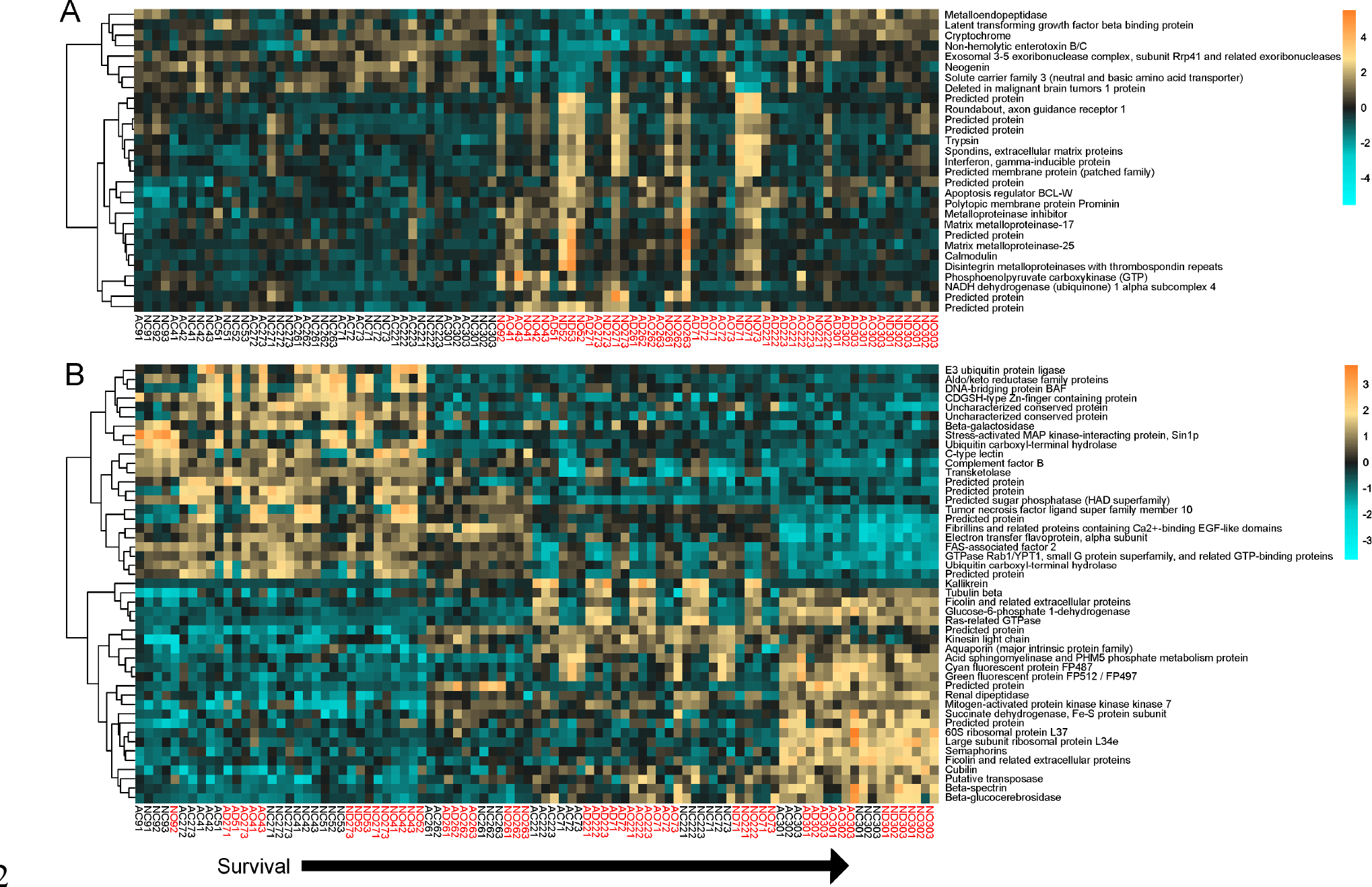
Response gene expression heatmaps. (**A**) Heatmap comparing the top DEGs (FDR = 1e-4) from response gene expression profiles by treatment. Samples are ordered by treatment, then by increasing survival from left to right. (**B**) Heatmap comparing the top DEGs (FDR = 1e-9) from response gene expression profiles by survival fraction. Samples are ordered by increasing survival from left to right. Red text denotes treated samples. Columns, rows, clustering, and color scale as indicated in Fig. S2.

**Fig. S4:**
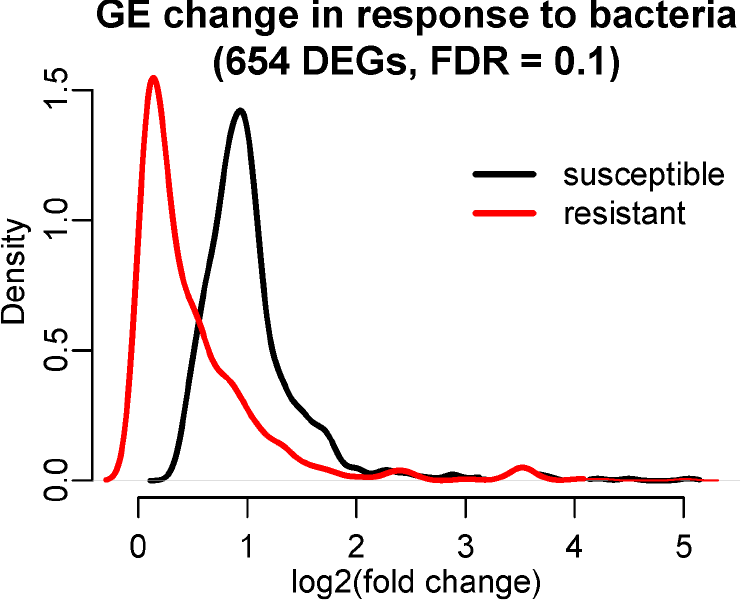
Density plots of predicted gene expression changes in response to bacteria treatment in fully susceptible corals (survival = 0, black line) and fully resistant corals (survival = 1, red line). Expression changes are extrapolated from the DESeq2 model incorporating survival as a continuous predictor variable. The plots are based on 654 bacteria-responding DEGs identified at FDR = 0.1.

**Fig. S5:**
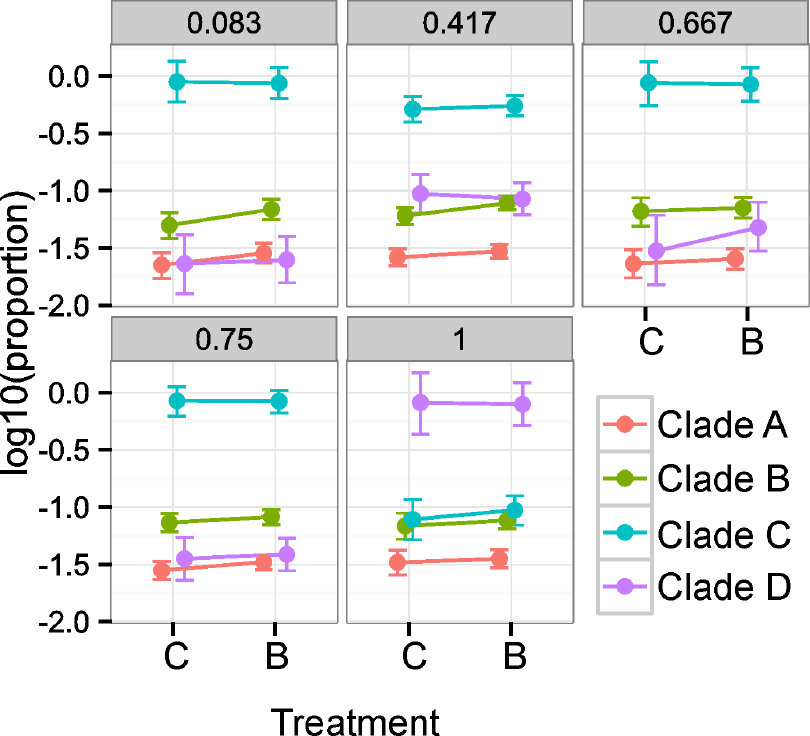
Relative abundances of *Symbiodinium* clades in *A. millepora* based on RNA-seq reads mapping to clade A, B, C, or D transcriptomes. The plots show posterior means and 95% credible intervals. Genotypes W30 and W27 were dominated by clade D. All other genotypes were dominated by clade C.

**Fig. S6:**
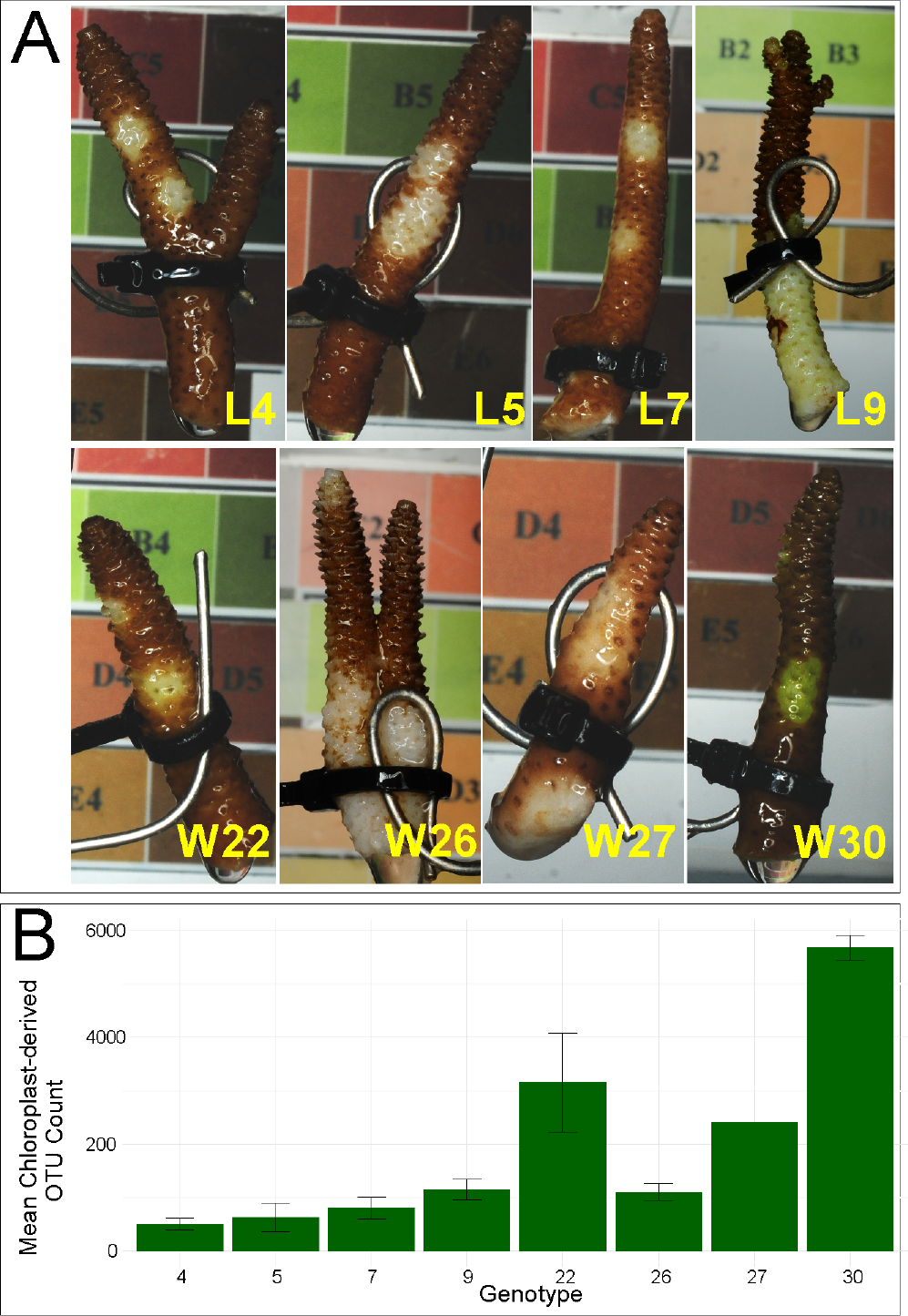
Example photos of each genotype and chloroplast-derived OTU counts. (**A**) Genotypes are indicated in yellow text. L4, L5, L7, W22, W27, and W30 were abraded with an airgun. Genotypes L9 and W26 demonstrate lesion development at ~50% tissue loss (time of death in this experiment). (**B**) Mean chloroplast-derived OTU counts (±SE) for each genotype

**Fig. S7:**
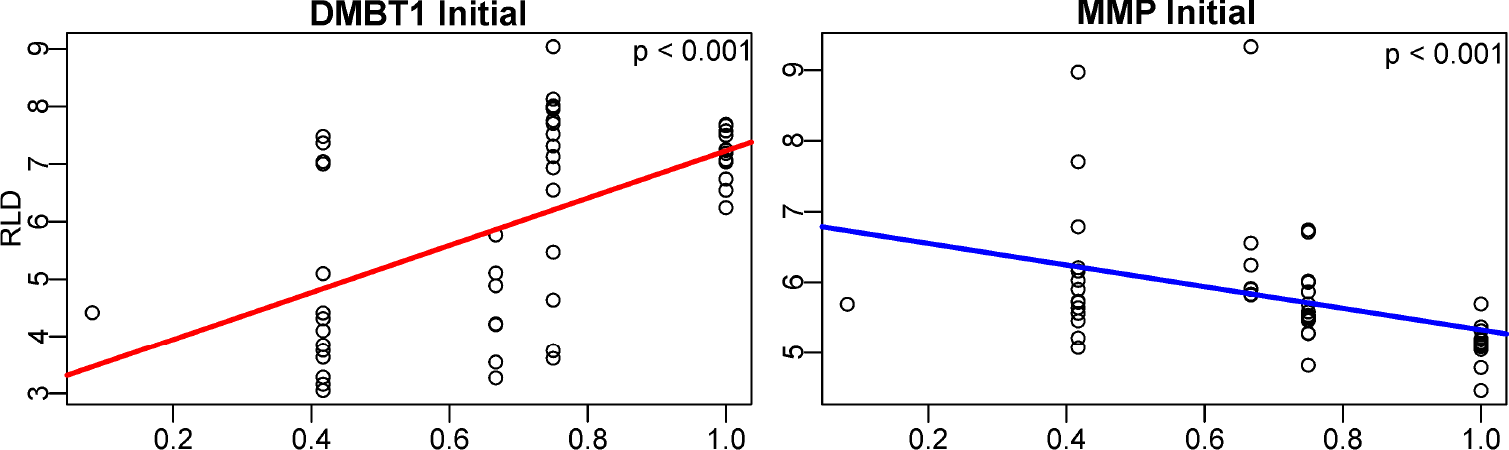
Initial biomarker expression. Initial correlations of rlog expression data (RLD) of *dmbt1* and *mmp* with survival fraction. P-values were generated by linear models testing the effect of survival.

**Table S1:** OTU group significance resu ts comparing bacteria-treated and control samples. Significant differences in OTU abundances between treatments were assessed using a likelihood ratio test as implemented by the G-test in QIIME. Taxonomy key: k = kingdom; p = phylum; c = class; o = order; f = family; g = genus; s = species.

| FDR | Treated | Control | Taxonomy |
| --- | --- | --- | --- |
| 0 | 61.86666667 | 0.0625 | k__Bacteria; p__Proteobacteria; c__Gammaproteobacteria; o__Vibrionales; f__Pseudoalteromonadaceae |
| 0 | 85.46666667 | 1.1875 | k__Bacteria; p__Proteobacteria; c__Gammaproteobacteria; o__Alteromonadales; f__Alteromonadaceae; g__Alteromonas |
| 0 | 114.7333333 | 10.5625 | k__Bacteria; p__Proteobacteria; c__Gammaproteobacteria; o__Alteromonadales; f__Alteromonadaceae |
| 8.27E-09 | 41.33333333 | 2.0625 | k__Bacteria; p__Proteobacteria; c__Gammaproteobacteria; o__Oceanospirillales; f__Oceanospirillaceae; g__Oleibacter |
| 9.31E-09 | 39.66666667 | 1.75 | k__Bacteria; p__Proteobacteria; c__Alphaproteobacteria; o__Rhodobacterales; f__Rhodobacteraceae |
| 5.07E-08 | 36.13333333 | 1.5625 | k__Bacteria; p__Proteobacteria; c__Alphaproteobacteria; o__Rhodobacterales; f__Rhodobacteraceae |
| 0.00012264 | 18.6 | 0.1875 | k__Bacteria; p__Proteobacteria; c__Gammaproteobacteria; o__Oceanospirillales; f__Oceanospirillaceae |
| 0.000542149 | 15.06666667 | 0 | k__Bacteria; p__Proteobacteria; c__Gammaproteobacteria; o__Vibrionales; f__Vibrionaceae; g__Photobacterium |
| 0.015230429 | 10.33333333 | 0 | k__Bacteria; p__Bacteroidetes; c__[Saprospirae]; o__[Saprospirales]; f__Saprospiraceae |
| 0.017515086 | 10.46666667 | 0.0625 | k__Bacteria; p__Proteobacteria; c__Gammaproteobacteria; o__Vibrionales; f__Vibrionaceae; g__Vibrio |
| 0.019538437 | 11.66666667 | 0.375 | k__Bacteria; p__Proteobacteria; c__Gammaproteobacteria; o__Oceanospirillales; f__Oceanospirillaceae; g__Oleibacter |
| 0.047846309 | 8.4 | 0 | k__Bacteria; p__Proteobacteria; c__Gammaproteobacteria; o__Vibrionales; f__Vibrionaceae; g__Vibrio |
| 0.080346846 | 7.6 | 0 | k__Bacteria; p__Bacteroidetes; c__[Saprospirae]; o__[Saprospirales]; f__Saprospiraceae; g__Saprospira |

**Table S2:** **Microbe classifications.**

| Family | Type | Reference |
| --- | --- | --- |
| Dermabacteraceae | Health | (1) |
| Simkaniaceae | Health | (2) |
| Hyphomicrobiaceae | Health | (3) |
| Rhizobiaceae | Health | (4, 5) |
| Hyphomonadaceae | Health | (6) |
| Erythrobacteraceae | Health | (6, 7) |
| Sphingomonadaceae | Health | (3) |
| Alcaligenaceae | Health | (5) |
| Endozoicimonaceae | Health | (2, 8, 9) |
| Moraxellaceae | Health | (4, 5) |
| Xanthomonadaceae | Health | (3, 10) |
| Cellulomonadaceae | Stress | (11) |
| Corynebacteriaceae | Stress | (11) |
| Micrococcaceae | Stress | (12) |
| Saprospiraceae | Stress | (13) |
| Bacteroidaceae | Stress | (11) |
| Amoebophilaceae | Stress | (14) |
| Cytophagaceae | Stress | (15) |
| Flammeovirgaceae | Stress | (11, 13, 16) |
| Flavobacteriaceae | Stress | (11, 13, 16) |
| Bacillaceae | Stress | (11) |
| Clostridiaceae | Stress | (11) |
| Lachnospiraceae | Stress | (11) |
| Peptostreptococcaceae | Stress | (11) |
| Planctomycetaceae | Stress | (16) |
| Kordiimonadaceae | Stress | (14) |
| Phyllobacteriaceae | Stress | (17, 18) |
| Rhodobacteraceae | Stress | (13, 16) |
| Rhodospirillaceae | Stress | (15, 18) |
| Rickettsiaceae | Stress | (19) |
| Burkholderiaceae | Stress | (16) |
| Comamonadaceae | Stress | (11) |
| Neisseriaceae | Stress | (20) |
| Rhodocyclaceae | Stress | (11) |
| Bacteriovoracaceae | Stress | (15) |
| Desulfobulbaceae | Stress | (21) |
| Desulfovibrionaceae | Stress | (22) |
| Campylobacteraceae | Stress | (16) |
| Alteromonadaceae | Stress | (11, 14) |
| Colwelliaceae | Stress | (13) |
| Enterobacteriaceae | Stress | (11, 16) |
| Halomonadaceae | Stress | (11) |
| Oceanospirillaceae | Stress | (11) |
| Pseudomonadaceae | Stress | (11) |
| Piscirickettsiaceae | Stress | (11, 14) |
| Pseudoalteromonadaceae | Stress | (11) |
| Vibrionaceae | Stress | (10, 11, 23, 24) |

## Supporting Information

### Results

Sequencing yielded an average of 2,042,678 reads per sample. An average of 42.2% of these remained after filtering. Mapping efficiency averaged 67.3%. A total of 44,687 isogroups (clusters of contigs representing the same gene, from here on referred to as “genes”) were detected after mapping to the transcriptome. Reads were converted to “unique transcript counts” (UTCs) by removing PCR duplicates, yielding an average of 360,642 counts per sample. The expression dataset was subset for genes with a mean UTC greater than three, yielding 16,781 genes. No outlier samples were detected.

### Materials & Methods

**Coral collection.** *Acropora millepora* were sampled from 3-6 m depth at the lagoon at Lizard Island (14°41’13.64S:145°27.75E) and the sheltered side of Wilkie Island (13°46’43.33S:143°38.75E; n = 4 colonies per reef) in the Great Barrier Reef in October 2013. Colonies were maintained in an outdoor raceway under flow through conditions and filtered natural light until fragmentation in November 2013. Tissue samples for baseline gene expression and physiological traits were preserved in liquid nitrogen and colonies were further fragmented into nubbins (4-5 cm) and secured on wire hooks (n = 18 per genotype). Corals in the experiment were collected under the permit number G12/35236.1 by the Great Barrier Reef National Park Authority of Australia.

**Bacterial culturing.** *Vibrio owensii* strain DY05 and *V. diazotrophicus* were used in this study. *V. owensii* has been implicated as the pathogenic agent of a tissue-loss disease in a Hawaiian coral, *Montipora capita* (1). At the time of this study *V. diazotrophicus,* a nitrogen-fixing commensal member of the coral holobiont (2), had not been implicated as a causative agent of any coral disease. Single isolates of each bacterial strain were recovered from glycerol stocks on Difco Marine Agar-2216 (BD, Franklin Lakes, NJ, USA) at 28°C. Cultures were incubated overnight at 28°C with shaking (150 rpm) in Difco Marine Broth-2216 (BD). Overnight cultures were triple-washed in 0.04 μM-filtered seawater (FSW) by centrifugation at 5000 *g* for ten minutes and resuspended in FSW. Washed cells were diluted to a final concentration of 1 × 10^9^ colony forming units (CFUs) · mL^−1^ in FSW. Cell densities were determined by counting CFUs resulting from plated serial dilutions and constructing a cell density calibration curve of absorbance (595 nm) versus CFU number.

**Experimental aquaria and abrasion procedure.** Coral fragments were secured upright in individual jars containing 200 mL FSW. Fluorescent lights provided light on a 12:12 h day/night schedule and the temperature was maintained at 26-27°C. Marine agar plates containing single colonies of *V. owensii* and *V. diazotrophicus* were maintained at 28°C for the duration of the bacterial challenge experiment. Fresh cultures were prepared from single isolates daily per the inoculation and wash procedures described above. Triple-washed cultures were diluted to achieve a final cell density of 10^7^ CFU · mL^−1^. Half of the 18 coral fragments received two small (~1 cm^2^) abrasions with a high-pressure airgun. These small abrasions mimicked clean lesions associated injuries that occur in nature (*e.g.,* corallivorous fish bites).

**Enzymatic assays.** Tissue was removed from skeletons using an airbrush and cold extraction buffer (100 mM TRIS, pH 7.8, with 0.05 mM dithiothreitol). Airbrushed tissue slurries were homogenized with 1 mm glass beads (BioSpec, Bartlesville, OK, USA) by vortexing for two minutes. The tissue slurry was centrifuged at 4°C for 10 minutes at 3200 *g* to separate coral and algal fractions. The coral protein supernatant was removed and stored at-80°C until use. Surface area determinations of airbrushed skeletons were made following a modified wax dipping protocol (3). A standard curve was prepared from a series cylinders of known surface area dipped in paraffin wax (Gulf Wax, Roswell, GA, USA) at 60°C. Coral skeletons were weighed and dipped twice in paraffin wax (59-60°C), weighing after each wax dip. For each fragment, the difference between initial weight and weight after second dip was compared to the standard curve to yield surface area in cm^2^. Protein measurements and enzymatic assays were performed using previously established protocols for coral immune studies (4–6).

Total protein was assessed in triplicate using the RED660 protein assay (G Biosciences, St. Louis, MO, USA) with a standard curve prepared from bovine serum albumin. Sample absorbance at 660 nm was compared to the curve and normalized to surface area and the tissue slurry volume.

Prophenoloxidase (PPO) activity was assayed in triplicate by mixing 20 μL of sodium phosphate buffer (50 mM, pH 7.0), 25 μL of trypsin (0.1 mg · mL^−1^), and 20 μL of protein extract. Dopamine (30 μL, 10 mM) was added as substrate and absorbance at 490 nm was measured every 30 seconds for 15 minutes. Change in absorbance was calculated during the linear range of the curve (1-3 minutes). Activity was expressed as the change in absorbance per mg of protein (ΔA_490_ · mg protein^−1^ · min^−1^). Phenoloxidase (PO) activity was assayed in triplicate by mixing 20 μL of sodium phosphate buffer (50 mM, pH 7.0), 25 μL of sterile water, and 20 μL of protein extract. Dopamine (30 μL, 10 mM) was added as substrate and absorbance at 490 nm was measured every 30 seconds for 15 minutes. Change in absorbance was calculated during the linear range of the curve (1-3 minutes). Activity was expressed as the change in absorbance per mg of protein (ΔA_490_ · mg protein^−1^ · min^−1^). Catalase (CAT) activity was assayed in triplicate by mixing 45 μL of sodium phosphate buffer (50 mM, pH 7.0), 75 μL of 25 mM H_2_O_2_, and 5 μL of protein extract. Samples were loaded on ultraviolet transparent plates (UltraCruz, Santa Cruz Biotechnology, Dallas, TX, USA) and absorbance at 240 nm was measured every 30 seconds for 15 minutes. Change in absorbance was calculated during the linear range of the curve (1-3 minutes). Activity was expressed as the change in hydrogen peroxide concentration per mg of protein (ΔH_2_O_2_ · mg protein^−1^ · min^−1^). Peroxidase (POX) activity was assayed in triplicate by mixing 40 μL of sodium phosphate buffer (10 mM, pH 6.0), 25 μL of 10 mM guaiacol, and 10 μL of protein extract. Absorbance at 470 nm was measured every 30 seconds for 15 minutes. Change in absorbance was calculated during the linear range of the curve (1-3 minutes). Activity was expressed as the change in absorbance per mg of protein (ΔA_470_ · mg protein^−1^ · min^−1^).

**Gene expression library preparation.** RNA was extracted (RNAqueous Total Isolation Kit, Life Technologies, Carlsbad, CA, USA), treated with DNase I (Life Technologies), and heat-fragmented. Then it was converted to PCR-amplifiable first-strand cDNA using oligo-dT-containing primer, a template-switching oligo, and SMARTScribe Reverse Transcriptase (Clontech, Mountain View, CA, USA). The libraries were PCR-amplified, individually barcoded, size-selected, pooled, and sequenced on the Illumina HiSeq platform (version 2500) at UT Austin’s Genome Sequencing and Analysis Facility.

**Gene expression analysis.** The reads were trimmed, deduplicated, quality filtered, mapped to the *A. millepora* reference transcriptome (7) using *bowtie2* (8), and converted to UTCs representing the number of independent observations of a transcript of a specific gene, summed over all isoforms. All subsequent analyses were carried out in R version 3.1.3 (9). Sample outliers were detected using *arrayQualityMetrics* (10). Low-expressed genes with a mean UTC less than three across all samples were discarded from the analysis.

**Primer design.** For *deleted in malignant brain tumors protein 1,* the forward and reverse primers were 5’-TCATGTGACCTGTGTTGGGA-3’ and 5’-GGTGACGCTCCGATCAAAC-3’, respectively. For *gelatinase A and related matrix metalloproteases*, the forward and reverse primers were 5’-GTTCCAAAATCGGCCACACC-3’ and 5’-CGTTATGCAGGGCTTCCAGA-3’, respectively. Primer pair specificity was verified by gel electrophoresis and melt curve analysis of the amplification product obtained with template *A. millepora* cDNA. Primer efficiencies were determined by amplifying a series of two-fold dilutions of *A. millepora* cDNA and analyzing the results using function *PrimEff* of the *MCMC.qpcr* package in R (11). Briefly, C_T_ (threshold cycle) results were plotted as C_T_ vs. log_2_[cDNA], and amplification efficiencies (amplification factor per cycle) of each primer pair were derived from the slope of the regression using formula: efficiency = 2^−(1/slope)^ (12).

**Microbiome community analysis.** DNA was isolated using an RNAqueous kit together with the RNA for gene expression analysis. DNA samples were diluted to 10 ng · μL^−1^. The bacterial 16S rRNA gene V4/V5 region was amplified using the Hyb515F (5′-TCGTCGGCAGCGTCAGATGTGTATAAGAGACAGGTGYCAGCMGCCGCGGTA-3′) and Hyb806R (3′-TAATCTWTGGGVHCATCAGGGACAGAGAATATGTGTAGAGGCTCGGGTGCTC TG-5′) primers and sequenced on the MiSeq V2 platform to generate 250 bp paired reads. Sequences with six or more consecutive identical bases (12,128 sequences) or incorrect primer sequence (63,719 sequences) were discarded using split_libraries.py in QIIME (Quantitative Insights Into Microbial Ecology (13)). Sequences of 97% similarity were clustered into operational taxonomic units (OTUs). A phylogeny was generated by aligning representative sequences that were filtered to remove gaps and hypervariable regions.

